# Ongoing repair of migration-coupled DNA damage allows stem cells to reach wound sites

**DOI:** 10.1101/746701

**Authors:** Sounak Sahu, Divya Sridhar, Prasad Abnave, Nobuyoshi Kosaka, Anish Dattani, James M. Thompson, Mark A. Hill, A. Aziz Aboobaker

## Abstract

The impact of mechanical stress during cell migration may be a previously unappreciated source of genome instability [1–3], but to what extent this happens *in vivo* remains unknown. Here we consider an *in vivo* system where the adult stem cells of planarian flatworms are required to migrate to a distal wound site [4]. We observe a relationship between adult stem cell migration and ongoing DNA damage and repair during tissue regeneration. Migrating planarian stem cells undergo changes in nuclear shape and increased levels of DNA damage. Increased DNA damage levels resolve once stem cells reach the wound site and stop migrating. Stem cells in which DNA damage is induced prior to wounding take longer to initiate migration suggesting migration activity is sensitive to DNA damage. Migrating stem cells populations are more sensitive to further DNA damage than stationary stem cells, providing evidence that levels of migration-coupled-DNA-damage (MCDD) are significant. RNAi mediated knockdown of DNA repair pathway components blocks normal stem cell migration, confirming that DNA repair pathways are required to allow successful migration to a distal wound site. Together these lines of evidence demonstrate that migration leans to DNA damage in vivo and requires DNA repair mechanisms. Our findings reveal that migration of stem cells represents an unappreciated source of damage, that could be a significant source of mutations in animals during development or during long term tissue homeostasis.

## Introduction

Both exogenous and endogenous agents constantly threaten genome integrity and cells have evolved mechanisms to counteract these various forms of DNA damage [5,6]. Accumulated genotoxic damage, particularly in stem cells, is thought to underpin both ageing and the development of cancer [7–10]. Therefore, maintaining the integrity of the genome is particularly important for longer-lived animals with adult stem cells. We know that efficient DNA repair mechanisms to counteract the effects of replicative stress and other sources of damage are central to avoiding both premature ageing and cancer [11–14]. Recent *in vitro* studies using cancer cell lines and dendritic cells have shown that migration through micro capillaries imparts mechanical stresses on nuclei, and this can be a source of genome instability [1,2,15–17]. If migrating cells in vivo, in particular stem cells, experience genome instability this has a number of broad implications for metazoan development and homeostasis. If migrating cell populations experience more DNA damage they maybe more likely to be become transformed to cause malignancies, or become senescent and contribute to ageing.

However, it is not known to what extent the is important *in vivo* in adult stem cells. To address this, we have studied DNA damage and repair processes in the highly regenerative planarian *Schmidtea mediterranea*, and in particular looked to see if they have a role in migrating adult stem cells.

## Materials and Methods

### Planarian culture

Asexual freshwater planarians of the species *Schmidtea mediterranea* were used in this study. The culture was maintained in 0.5% Instant Ocean and fed with organic calf liver twice a week. Planarians were starved for 7 days prior to each experiment and also throughout the duration of each experiment and cultured in the dark at 20°C.

### Gene cloning and RNAi

The sequences of S. mediterranea RAD51 and BRCA2 were described previously. Planarian DDR genes were identified by BLAST searches against the the Planmine database, leading to the identification of full-length mRNA transcripts (Smed-ATR-dd_Smed_v6_8754_0_1, Smed PARP1-dd_Smed_v6_10338_0_1, Smed PARP2-dd_Smed_v6_6154_0_1, Smed PARP3-dd_Smed_v6_2611_0_1, Smed-FANCJ-dd_Smed_v6_16638_0_2). Fragments of these genes were cloned into the pPR-T4P plasmid vector containing opposable T7 promoters (kind gift from Jochen Rink, MPI Dresden). These clones were used for in vitro transcription to synthesize dsRNA and probes as previously described [4]. dsRNA was delivered via microinjection using Nanoject II apparatus (Drummond Scientific) with 3.5” Drummond Scientific (Harvard Apparatus) glass capillaries pulled into fine needles on a Flaming/Brown Micropipette Puller (Patterson Scientific). Worms were injected with 3×32nl of dsRNA 6 times over 2 weeks. A 1-day gap was kept between the last injection and Irradiation experiments (as described in [4]).

Primers used for amplification of DNA for dsRNA synthesis/ RNA probes can be found in **Supplementary Table 1**.

### Gamma irradiation

Animals were starved for at least 7 days and exposed to 1.5,5,10,15,20 and, 30 Gy of ^137^Cs gamma-rays (for Fig.1 A) using a GSR D1 Gsm (Gamma-Service Medical GmbH, Leipzig, Germany) gamma irradiator at a dose rate of 1.9 Gy/min.

**Figure 1.**
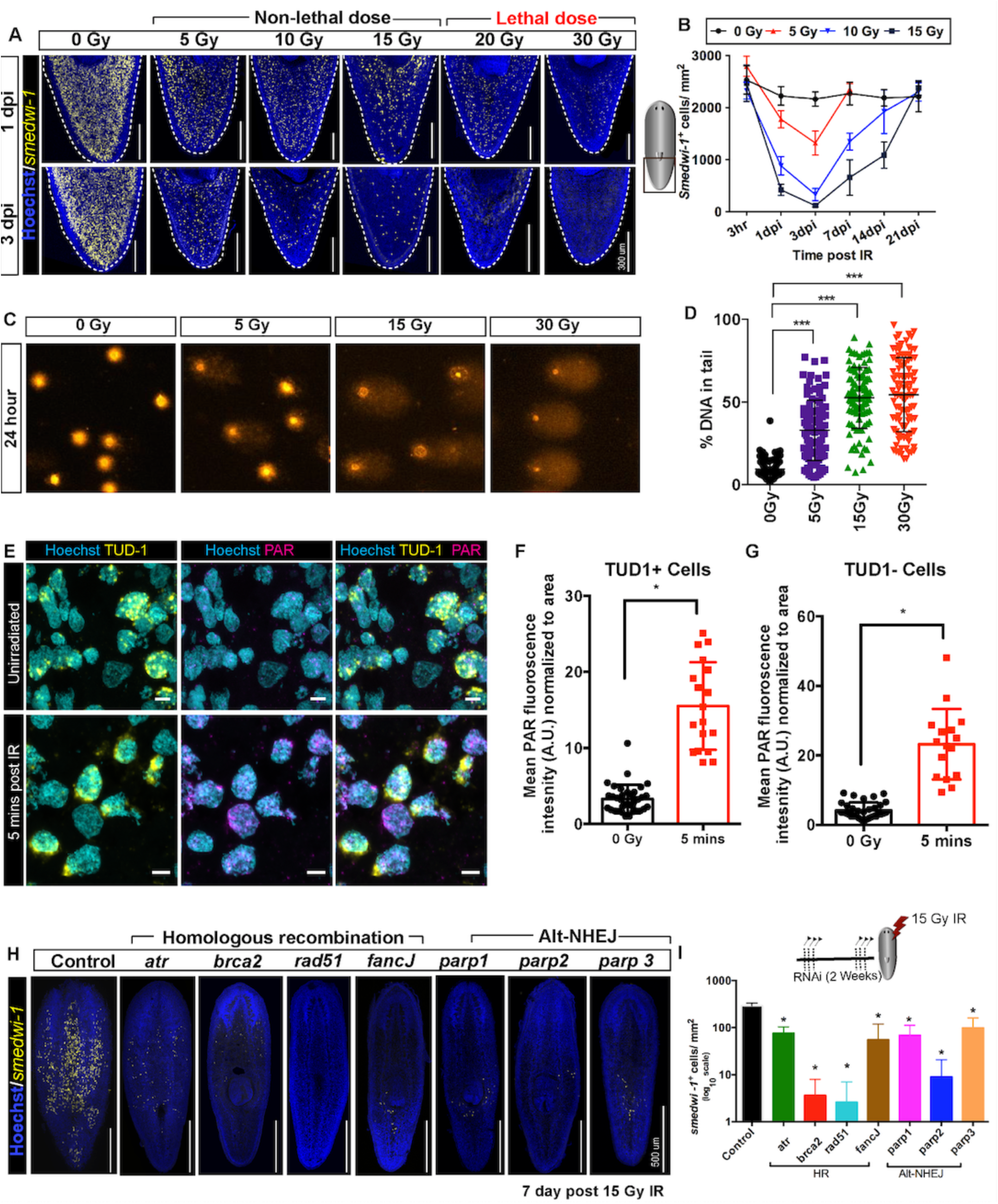
Planarian stem cell resistance to doses up to 15 Gy of gamma IR requires conserved DDR pathways. **(A)** *smedwi-1* FISH of planarians exposed to different doses of gamma IR (5,10,15,20 and 30 Gy) after 1 and 3 days post IR (dpi) showing a dose dependent decrease in stem cell number. Scale bar: 300 μm. **(B)** Quantification of *smedwi-1*^*+*^ cells/mm^2^ (yellow) showing the repopulation kinetics of surviving stem cells after different doses of IR post IR (n = 5 per dose, per time point). Results are expressed as mean ± SD. **(C)** COMET assay showing the extent of DNA breaks (comet shape) in isolated planarian cells at 24 hours after exposure to 5, 15 and 30 Gy of IR. **(D)** Quantification of the percentage of tail DNA in COMET assay post IR at 24 hours. Results are expressed as mean ± SD. Each dot represents the tail DNA in individual planarian cells. **(E)** Double immunostaining with Anti Tudor-1(Yellow) and Anti-PAR (Magenta) showing DNA damage in stem cells (Tud1^+^) and post mitotic differentiated cells (Tud1^−^) at 5 mins post 5 Gy IR. Nucleus is stained with Hoechst (Blue). **(F-G)** Quantification of PAR fluorescence in Tud1^+^ and Tud1^−^ cells normalized to the nuclear area in irradiated and unirradiated cells. **(H)** Representative FISH showing stem cell repopulation in Control (gfp) RNAi and after knockdown of different DNA repair genes [involved in homologous recombination (*atr, brca2, fancJ, rad51*) and Alt-NHEJ (*parp1, parp2, parp3*)] after 7 days post 15 Gy IR. **(I)** Repopulation of *smedwi-1*^*+*^ cells/mm^2^ in DDR RNAi worms after 7 days post 15 Gy IR (n = 5 per condition). Results are expressed as mean ± SD in log_10_ scale (student’s t-test; *p<0.05).

### Shielded Irradiation assay

Shielded irradiation was performed as previously described [4]. Worms (3-5mm) were anesthetized in ice-cold 0.2% chloretone and aligned on 60 mm petridish. The petridish was pre-marked with a line at the bottom according to the dimension of the shield (as described in Supplementary Fig 5A). The anterior tip of individual worms were aligned to keep the absolute migratory distance between tip of head and shielded region fixed and the petri dish containing worms was then placed directly on top of the lead shield and irradiated from below with X-rays (225kV, 0.5mm AI filter, 23 Gy/min). The head and tail regions of the worms received 30 Gy while the shielded region received less than 1.5 Gy (Supplementary Fig 5B). Immediately following irradiation the worms were placed into fresh planarian water and cultured in the dark at 20°C. For experiments involving an initial dose of Gamma IR, worms are incubated for 15 mins in planarian water before used for shielded irradiation assay (Experiment in Figure 3 A). Heads were amputated 4 days post shielded irradiation to induce migration towards the wound (considered as 0 day post amputation (dpa) or modified as necessary for a particular experiment (e.g. Figure 3A). Lack of posterior cell migration in the absence of a posterior wound allowed us to define the boundary of the shield and measure the distance migrated by stem cells in whole mount samples as previously described [4].

### COMET Assay in planarians

Frosted microscope slides were coated with 700μl of 1% Normal Melting Point Agarose (NMPA) in 1X PBS to make a uniform layer and dried overnight at 55°C. Worm fragments were gently diced to reduce any mechanical stress to the cells. The tissue pieces were digested using Papain (Sigma Aldrich) (15 U/ml) for 1 hour at 25°C. Pieces were mechanically dissociated using a P1000 pipette to form a single cell suspension and filtered through 100 μm and then 35 μm cell strainers (BD Falcon). 10,000 dissociated single cells were re-suspended in 80μl of CMFHE^2+^ and an equal amount of 1.5% Low Melting Point Agarose added and mixed. 40 μl of the cell-agarose suspension was added onto an NMPA coated slide and allowed to solidify at 4°C. Slides were incubated overnight (~15 hours) in a coplin jar at 4°C with 89% Lysing solution [(2.5 M Nacl, 100mM EDTA, 10 mM Trizma base, NaOH added to pH 10.0) and freshly added 1% Triton x-100]. This solution was then replaced with neutralization buffer (0.4M Tris base in dH_2_O, pH to 7.5) for 15 mins at 4°C. The neutralization buffer was then removed, and slides were placed into an electrophoresis chamber at 4°C filled with freshly prepared 1X electrophoresis buffer (300mM NaOH and 1mM EDTA in dH_2_O). The slides were allowed to equilibrate for 15 minutes followed by an alkaline electrophoresis at 20 V for 20 minutes at 4°C. Next, slides were transferred back into the coplin jar and equilibrated for 5 minutes in neutralization buffer. The slides were stained with SYBR Green I (1:10000 dilution) in freshly prepared 1X TE buffer (10mM Tris-HCl and 1mM EDTA, pH 7.5). For long term storage the slides were fixed with cold 100% ethanol for 5 minutes and dried. After drying 50 comets per slides were analysed using KOMET software (Andor) and the % of tail DNA was measured after different doses of gamma irradiation, or from “shielded” regions or “migrating” regions both with and without migrating stem cells (i.e. with our without wounding).

### *In situ* hybridization and Immunostaining

*In situ* hybridization was performed as described in [37]. Previously reported sequences were used for riboprobe synthesis of *smedwi-1* [4]. DIG-labelled RNA probes were generated for *atr, atm, brca2, parp-1, parp-2 and, parp-3* and co-localised using FITC labelled *smedwi-1* probe. Antibodies used for Immunostaining were anti-H3P (phosphorylated serine 10 on histone H3; Millipore; 09-797; 1:1000 dilution [4]), anti-Poly (ADP) Ribose [21] (PAR) monoclonal antibody (1:250) (Santacruz, clone 10H), anti-TUD1 (1:250 dilution, based on [39]). The secondary antibodies used were Anti-Rabbit-HRP and anti-mouse-HRP respectively (1:2000 dilution) and Tyramide signal amplification was performed for FISH and immunostaining as described in [37].

### Sectioning of planarian worms

Panarians were killed in 2% HCl and Holtfreter solution and fixed with 4% formaldehyde for 2 hours. The worms were then washed in PBST (0.3% Triton-X) and dehydrated with an increasing gradient of methanol washes and stored at −20°C. The following day worms were re-hydrated with a decreasing gradient of Methanol and PBS, Xylene washes (2 washes of 7 minutes each) and put into molten paraffin for 1 hour. Individual worms were then aligned (sagittal or transverse) in paraffin moulds, trimmed and sliced into 10 μm sections using a microtome. Individual ribbons of planarian sections were put in a 37°C water bath and aligned to have the entire worm on each Poly-lysine coated slide.

### Immunostaining on paraffinized sections

Planarian sections were deparaffinized using Xylene substitute (2 washes of 7 minutes each) and washed with PBS-Tx0.3 (0.3% Triton-X). The sections were subjected to antigen retrieval with Trilogy (Cell Marque) at 90°C. The slides were then fixed in 4% fixative for 15 mins followed by two washes of PBS-Tx0.5 (0.5% Triton-X) for 30 mins and transferred to a blocking solution (0.5% BSA and PBS-Tx0.5). 150 μl of primary antibody (diluted in blocking solution) was added to individual slides and a Parafilm was placed on top for uniform spreading of the antibody solution. After an overnight antibody incubation at 4°C, the slides were washed with alternating changes of PBS-Tx0.5 and PBS+0.1% Tween-20. The secondary antibodies were diluted in blocking solution and incubated overnight. Slides were washed again with alternating changes of PBS-Tx0.5 and PBS+0.1% Tween-20. After two 10 minutes washes slides were developed with Tyramide/other fluorophores. For double immunostaining, Sodium-Azide based peroxide inactivation was performed after the development of each antibody. After two 10 min washes with PBS-TW, slides were stained with DAPI for nuclear staining overnight. Slides were mounted and imaged in 100X oil objective lens in Olympus FV1000 confocal microscope with the appropriate fluorescent laser.

### Image processing and data analysis

Whole worm confocal imaging was done with Olympus FV1000 and taken as Z-stacks (slices of 4 μm each D/V axis) stitched and then processed as a maximum projection using Fiji software. All measurements and quantifications were done with Fiji https://fiji.sc/ (using cell counter plug in) and normalized to the area. Images in Figure 2 (A-B, i-iv) are single confocal stacks (0.32μm) taken in Zeiss 880 Airyscan microscope using 63X Oil objective lens and manually cropped into individual cells for counting in Fiji software. Nuclear aspect ratio is measured taking the ratio of the length and the width of the nucleus[33]. Images in Supplementary Figure 3 are single confocal stacks (0.5μm) taken in Olympus FV1000 using 100X oil objective lens. Images were processed/cropped/and signals pseudo-coloured in Fiji software. Background is changed into black for better visualisation and all figures are prepared using Adobe Illustrator v6. PAR fluorescence in Fig. 1 E-G, 2 D-F and Supp. Fig 2 D-F was performed in Fiji using the “mean fluorescence analyse” tool and normalized to the nuclear area using the Hoechst signal. Total PAR fluorescence was measured from all the nuclei, with thigh intensity of perinuclear TUD1 staining determining Tudor-1 positive stem cells [39].

**Figure 2.**
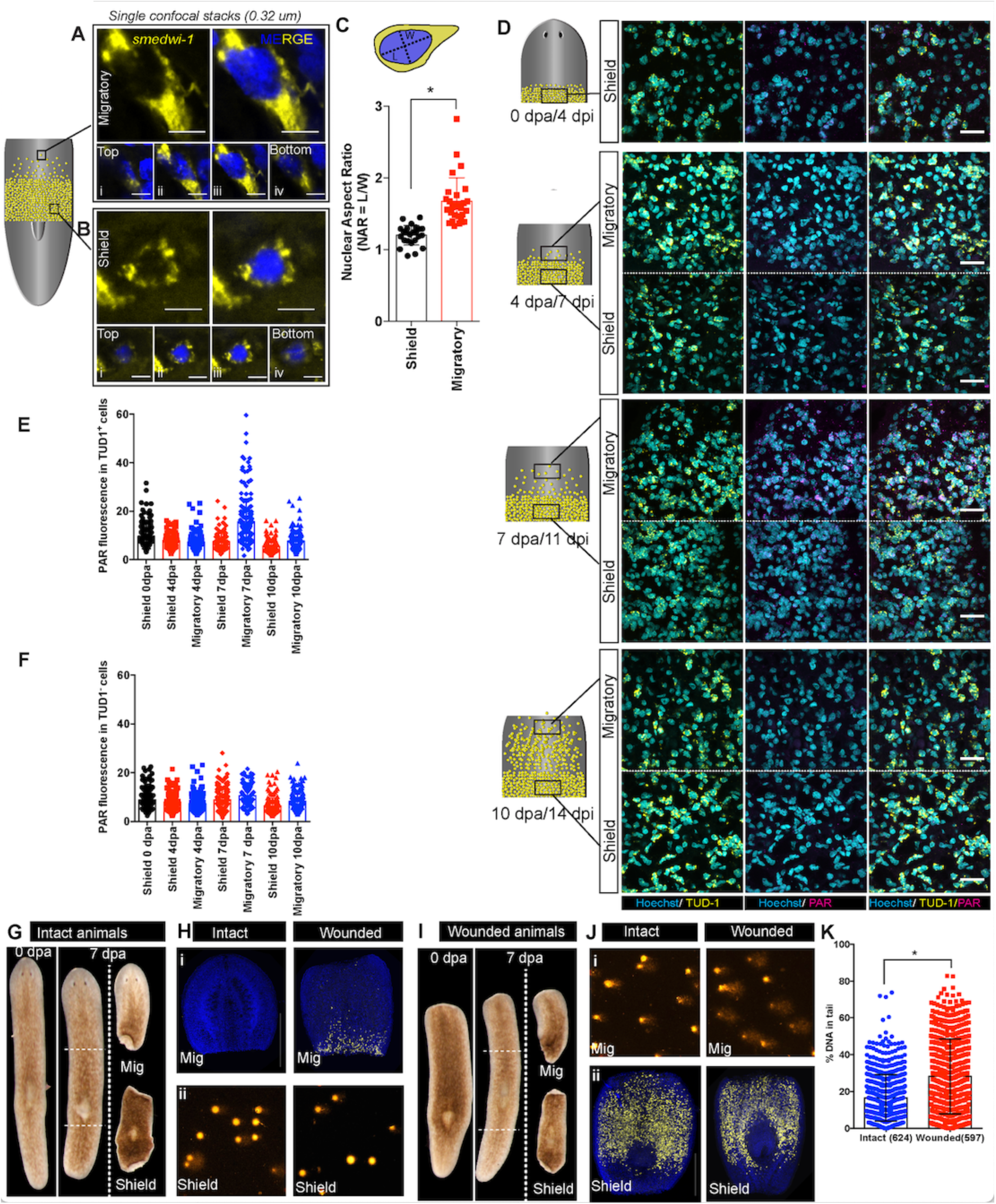
Migration-coupled DNA damage (MCDD) in stem cells. Representative FISH showing stem cells (*smedwi-1*^*+*^) with extended protrusions in migratory cells **(A)** and stationary cells from the shielded region **(B).** Nuclei stained with hoechst (blue). Images are shown as single confocal Z-stack (0.32μm). (i-iv) is the single Z-stack from top to bottom. Scale bar: 5 μm. **(C)** Quantification of Nuclear aspect ratio (NAR) in the migratory cells compared to stationary cells (n = 28 cells in migratory region and n = 24 cells from the shield at 7dpa/11 dpi (shielded irradiation assay) **(D)** Immunostaining with anti-PAR (magenta) in migrating stem cells (anti-TUD1, yellow) after 0, 4, 7 and 10 days post amputation showing MCDD in TUD1+ migrating stem cells. Box denotes the field of cells imaged for analysis. Nuclei stained with Hoechst (blue). Scale bar: 25 μm. Quantification showing the PAR fluorescence normalized to the nuclear area from Tud1+ stem cells **(E)** and post-mitotic differentiated Tud1-cells **(F)** in the migrating region compared to stem cells in the shield **(G, I)** Brightfield images of intact (G) and wounded (I) animals at 0 dpa and 7 dpa showing the amputated migratory region and shielded region. Dotted lines denote the position of the shield. The migratory region was used for *smedwi1* FISH and the corresponding shielded region was used for COMET assay and vice versa depending on the context (refer to Supp. Fig. 6). **(H)** *smedwi-1* FISH of the migratory tissues at 7dpa showing the presence of migrating stem cells in wounded animals compared to no migration in intact animals. Cells corresponding to the shielded region were used for COMET assay to check for the extent of DNA damage. **(J)** *smedwi-1* FISH of the shielded tissue at 7dpa showing the presence of stem cells under the shield in intact and wounded animals. Cells corresponding to the migratory region from the animals were used for COMET assay to check for the extent of DNA damage in migrating stem cells in wounded animals. **(K)** Quantification of COMET assay showing the extent of DNA breaks in migrating cells in wounded animals compared to intact animals (absence of migrating stem cells). Results are expressed as mean ± SD (student’s t-test; *p<0.05). Each dot represents the percentage of tail DNA from single cells after COMET assay (n=624 cells from intact animals, and 597 cells in wounded animals). [See also Supplementary Figures 6].

**Figure 3.**
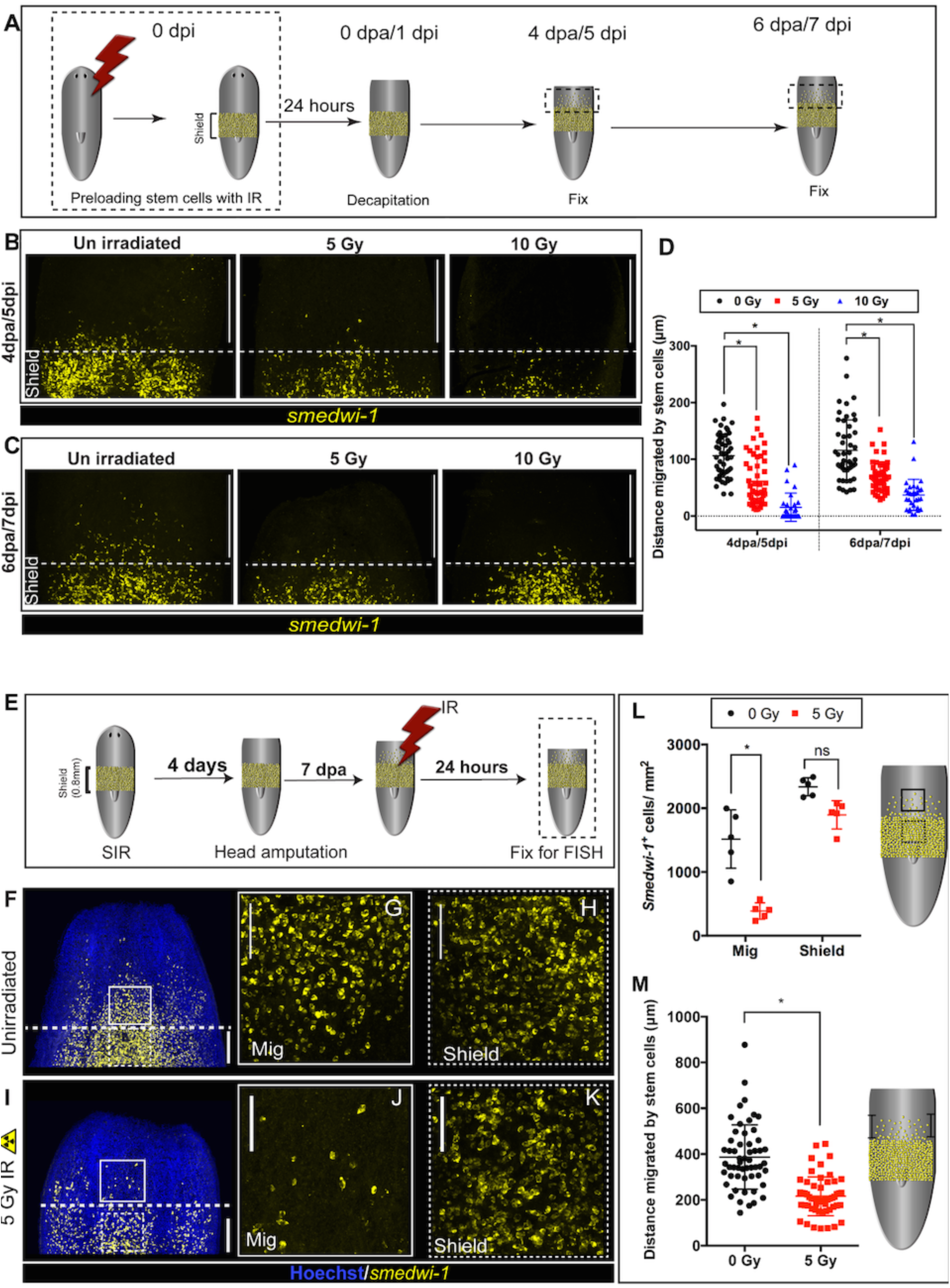
DNA damage delays migration and migrating stem cells with MCDD are more sensitive to ionising radiation. **(A)** Experimental scheme showing worms pre-exposed to irradiation (5 and 10 Gy) followed by a shielded irradiation and decapitation after 24 hours. Worms are fixed at 4dpa and 6dpa (dpa = days post amputation). Box represents the migratory region, represented in the figure below. **(B-C)** FISH showing worms pre-exposed to IR (5 and 10 Gy) show delayed stem cell migration after 4 and 6dpa. Dotted line represents anterior boundary of the shield. Scale bar: 350 μm **(D)** Distance migrated by 10 most distant cells are counted from individual worms (n = 5 per condition) Results are expressed as mean ± SD. Statistical significance determined by multiple t-test using the Holm-Sidak method, *p<0.05 **(E)** Schematic of experimental set up to study sensitivity of migrating cells to IR. In addition to the initial shielded irradiation, the worms were irradiated with a low dose of IR (5 Gy, whole body) when MCDD is high (7dpa) and are fixed after 24 hours to check the survival of the migratory stem cells to IR. **(F-K)** Representative *smedwi-1* FISH showing migrating cells are more sensitive to IR than the cells in the shielded region. The region counted for analysis is marked with a box (bold: migratory region, dotted: Shielded region). [n = 5 per condition, scale bar: 200 μm (F, I), 100 μm (G, H, J, K)]. **(L)** Quantification of *smedwi-1*^*+*^ cells/mm^2^ cells in the shielded region and in the migratory region. The decrease in cells/ mm^2^ in the migratory field is significant compared to the decrease in the shielded region indicating that MCDD sensitises cells to IR. Cartoon showing the region counted for analysis. Each dot represents number of surviving cells from individual worms, n = 5. Statistical significance determined by Tukey’s multiple comparison test (*p < 0.05). **(M)** Distance migrated by stem cells showing that cells are more sensitive to low dose IR the further they have migrated. Each dot represents the distance migrated by individual cells. Distance migrated by 11 most distant cells are counted from individual worms (n = 5 per condition) Results are expressed as mean ± SD (student’s t-test; *p<0.05, ns = not significant).

### Single cell expression data

These data were taken from https://radiant.wi.mit.edu/app/[38]

### Expression based proportional data

These data were extracted from Dattani et al. Genome Research, 2018 [25]

### Statistical analysis

Results are expressed as mean ± standard deviation (SD). Statistical analyses were performed using Prism, GraphPad Prism version 6.0 (https://www.graphpad.com/). Student’s t-test and Tukey’s multiple comparison test was used for statistical significance at p<0.05.

## Results and Discussion

### A robust DNA damage response allows stem cells to resist doses up to 15 Gy of ionising radiation

Planarians have a population of pluripotent adult stem cells, and potentially avoid both ageing and cancer [18–20]. While a lethal ionising radiation (IR) doses of 20 and 30 Gray (Gy) both leave some *smedwi1+* stem cells after 24 hours, none of these remaining cells are competent to proliferate and rescue the animal. However, enough cycling *smedwi1+* stem cells that survive a 15 Gy dose are competent to proliferate, so that all animals survive this dose. (Figure 1 A, B; Figure S1 A - D). Stem cell loss after sub-lethal IR is dose dependent and continues until 3 days post irradiation (pi) (Figure 1B)

We optimised the use of the COMET assay and staining with antibodies to poly-ADP ribose (PAR), the currently available methods for measuring DNA damage/repair in planarians [21]. We observe that damage assayed by single cell gel electrophoresis of whole planarian stem cell populations is IR dose dependent (Figure 1 C and D), with subsequent repair of surviving cells taking place over the subsequent 11 days (Figure S2 A, B). By combining PAR staining with an antibody to planarian Tudor-1 that marks the perinuclear RNP granules (chromatoid bodies) in *smedwi-1+* stem cells (Figure S2 C) we measured damage in stem cells and post-mitotic differentiated cells simultaneously. Levels of PAR staining in the nucleus increased by 5 minutes of exposure to 5 Gy IR (Figure 1 E-G) and returned to baseline 24 hours after exposure (Figure S2 D-F), indicating that this approach can measure levels of acute DNA damage and repair.

We identified conserved DDR genes from the *S. mediterranea* genome and transcriptome [22–24] (Figure S1 E) that are known to be essential elsewhere to repair DNA breaks. Using FACS-proportion based expression analysis [25], single cell expression data [26] and double Fluorescent In Situ Hybridisation (FISH) with the *smedwi-1* pan-stem cell marker, we found that transcripts of most DNA repair genes (*atr, atm, brca2, parp1*, and *parp2*) are enriched in stem cells (Figure S3). Only RNAi of *rad51* led to animal death. We found that RNAi of other conserved DDR genes individually (*atr, brca2, parp1, parp2, parp3)* during normal regeneration and homeostasis did not lead to any phenotypic defects in regenerating animals and did not affect stem cell number or proliferation over a time course of several weeks (Figure S4 A-H). Combinatorial RNAi of these DNA repair components, in pairs, did lead to significantly reduced stem cell numbers (Figure S4 I, J), demonstrating that knockdown effecting more than one pathway does lead to the loss of stem cells during normal tissue homeostasis. Whether these data reflect residual repair function due incomplete knockdown by RNAi, compensation between repair pathways or a mixture of the two remains unknown.

After sub-lethal IR exposure, surviving stem cells clonally expand to restore homeostatic and regenerative capacity in a dose dependent manner [20,27,28] (Figure 1 A, B, Figure S1 A-D). In this scenario RNAi of the individual conserved components of Homologous recombination (HR) (*atr, brca2, rad51, fancJ)* and alt-Non-Homologous End-joining pathways *(parp1, parp2 and parp3)* after 15 Gy IR led to a failure in stem cell repopulation and animal death (Figure 1H – I, Figure S4 A-H). Together these data provide proof of principle that well-known DDR genes have an ongoing role in stem cell survival, and in DNA repair after IR exposure, establishing a basis for using *S. mediterranea* as an experimentally tractable *in vivo* model for studying DDR in adult stem cells.

### Migrating stem cells undergo Migration-coupled DNA damage (MCDD) that resolves at the wound site

Planarian stem cells and their progeny must migrate to the site of a wound or during reproductive (asexual) fission to form a regenerative blastema [29,30]. Recent work has shown mechanical stress on the cell nucleus, through a variety of proposed mechanisms, can lead to genome instability during cell migration [1–3]. However, how important this is generally in vivo in animals is unknown. To study this phenomenon in planarians, despite the lack of live-cell imaging approaches, we established a robust assay for stem cell migration [4] This uses a lead shield to perform ‘shielded irradiation’, to leave a stripe of stem cells whose subsequent migration can be followed (Figure S5 A – E). Decapitation triggers anterior migration from the shielded strip of stem cells towards the wound (Figure S5 C) and a lack of posterior cell migration over the experimental time course allows us to clearly define the posterior and therefore the pre-migration anterior boundary of the shield. This system has already allowed the detailed study of stem cell migration *in vivo* [4].

Using this assay, we asked if normal stem cell migration *in vivo* is genotoxic. Planarian stem cells are characterized by large nuclear: cytoplasmic ratios like other animals stem cells [31,32], suggesting that the nucleus in migrating stem cells could encounter physical stress through deformation of normal nuclear shape. In order to check nuclear shape plasticity, we measured the nuclear aspect ratio (NAR) [33] of the cells in the migratory region compared to stationary cells at 7 day post amputation and observed significant changes in NAR (Figure 2 A – C; Figure S5 F - G).

We then performed anti-PAR/Tudor-1 double immunostaining [21], to test whether we could detect increased DNA damage levels above normal background levels in migrating stem cells. We found that migrating TUD1+ cells accumulate increased levels of acute DNA damage measured by PAR staining (Figure 2 D, E), that eventually return to baseline levels when stem cells reach the wound site, are distributed through the irradiated tissue and cease migrating (Figure 2 E). Stationary stem cells remaining in the shield did not have raised levels of PAR staining implicating cell migration as the cause (Figure 2 D, E). Similarly, post-mitotic TUD-ve cells had lower levels of detectable PAR than migrating stem cells (Figure 2 F).

While PAR staining measures an acute response to new damage, a COMET assay measures global levels of DNA breaks, including those in cells undergoing apoptosis. We performed COMET assay on cells in the shielded and migratory regions of the migration assay (Figure S6 A). We used both intact (no migration) and 7 days post amputation (wounded) animals where stem cell migration is induced allowing comparison of cell populations from the migratory regions with and without migrating stem cells (Figure 2 G-K, S6A). We used in situ hybridisation to *smedwi-1* in animal fragments not used for COMET to confirm the accuracy of separating the shielded and migratory regions (Figure 2 H, J and S6 B-G). These experiments revealed that levels of COMET were increased in migratory regions containing migrating stem cells compared to migratory regions from intact animals that are devoid of stem cells (Figure 2 K). These data suggest that migrating stem cells have higher levels of DNA damage.

These experiments to measure NAR, levels of nuclear PAR, and DNA damage together suggest planarian stem cells undergo migration couple DNA damage (MCDD) in vivo.

### Stem cells pre-loaded with DNA damage incur a delay in migration

In order to understand whether the presence of DNA damage acts to inhibit active stem cell migration, we pre-irradiated whole worms with 5 and 10 Gy IR before following stem cell migratory behaviour (Figure 3 A, Figure S7 A). We decapitated animals within 24 hours to trigger stem cell migration while DNA damage is still present (Figure S2). We observed that pre-irradiated stem cells undergo a significant delay in migration (Figure 3 B - D), (Figure 3 D). Although there is a significant delay in migration, stem cells eventually reach the wound, maintain normal stem cell number, and fuel normal regeneration (Figure S7 B). These data suggest a relationship between active migration and levels of DNA damage, with increased levels of damage inhibiting active migration.

### Stem cells with MCDD show more sensitivity to IR

We next examined whether stem cells with MCDD are more sensitive to ionising radiation. If this were the case it would demonstrate that the increased damage observed during migration is a significant load on the repair machinery. In addition to performing shielded irradiation, the worms were given an additional dose of 5 Gy to the whole animal at 7 days post amputation (dpa) (Figure 3 E), a time point when the highest number stem cells are actively migrating [4] and when the most cells with increased levels of MCDD, measured by levels of nuclear PAR are observed, (Figure 2 E). We observed a significant decrease in stem cell survival in the migratory region after 5 Gy IR compared to stationary stem cells in the shield (Figure 3 F - M), demonstrating that migrating stem cells with MCDD are more sensitive to IR than stationary cells (Figure 3 H). We also found that those cells that had migrated furthest, but not yet reached the wound site were the most sensitive to IR (Figure 3 M), suggesting that accumulation of MCDD may on average increase with migratory distance. We did not observe this striking difference earlier in the migratory process (Figure S7 C - J) when cells have just started migrating in response to wounding or when cells had already reached the wound site and migration was complete (Figure S7 K - P). This demonstrates that increased IR sensitivity correlates with the levels of DNA damage, which is dependent on the extent of active stem cell migration (Figure 2 D - E).

### Wound induced stem cell migration requires active DNA repair mechanisms to resolve ongoing MCDD

We next asked whether active DNA repair pathways are a functional requirement for continued stem cell migration. We therefore performed RNAi of specific DDR genes in the context of the stem cell migration assay (Figure 4 A). We observed less migration in *atr, atm, brca2* and *parp1* RNAi worms compared to control RNAi worms (Figure 4 B - I). We did not observe any significant difference in stem cell number in the shielded region (Figure 4 H), suggesting the knockdown of these genes did not affect normal stem cell turnover, as we previously observed (Figure S4 A-H). The significant reduction in the distance migrated by stem cells (Figure 4 I) supports a role for active ongoing DNA repair in maintaining genomic integrity during migration and allowing migration to proceed in the face of ongoing damage to the genome.

**Figure 4.**
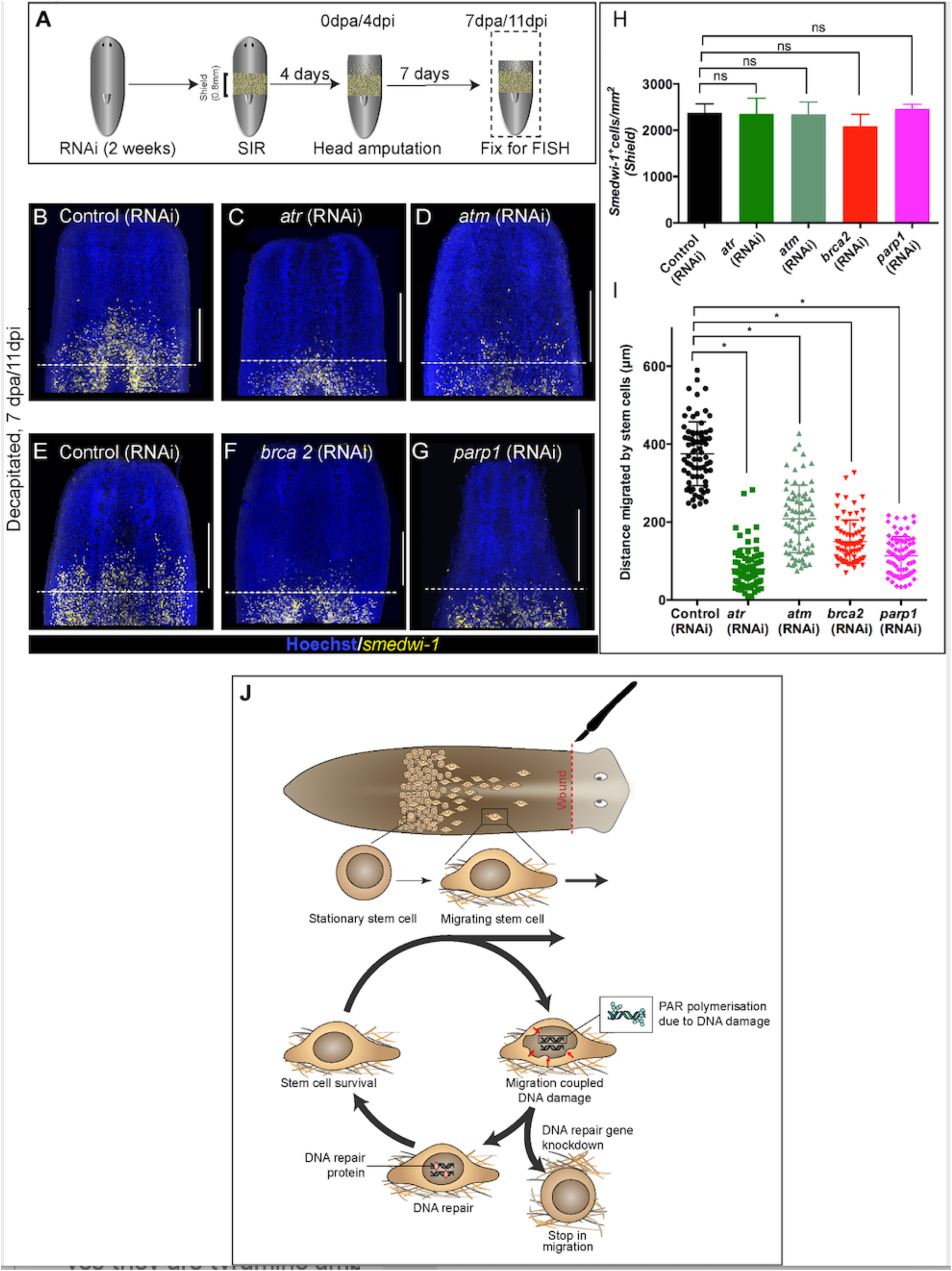
**(A)** Experimental scheme to study the role of DDR genes in stem cell migration. Worms are injected for 2 weeks (RNAi) followed by the shielded irradiation assay and fixed for FISH 7 days post head amputation. **(B-G)** Representative *smedwi-1* FISH shows migration of stem cells (yellow) at 7 dpa in control (RNAi) (B and E) worms, but migration is inhibited in *atr* (C), *atm* (D) *brca2* (F) and *parp1* (G) RNAi worms. (n = 5 per RNAi condition). Scale bar: 400 μm, dotted line represents the anterior boundary of the shielded region. **(H)** Stem cells in the shielded region show no significant changes in the stem cell turnover. (*p<0.05, ns= not significant, n=5 per RNAi condition). **(I)** Quantification showing the distance travelled by stem cells after knockdown of DNA repair genes compared to the control RNAi. Each dot represents the distance migrated by individual cells. Distance migrated by 15 most distant cells are counted from individual worms. Results are expressed as mean ± SD n=75 cells; N=5 worms/RNAi condition (student’s t-test; *p<0.05, ns = not significant). **(J)** Stem cells undergo changes in nuclear shape during migration compared to stationary cells in the shield. This model proposes that stem cells undergo migration, followed by MCDD and DNA repair. In the absence of functional DNA repair machinery stem cells fail to migrate.

## Conclusions

Here we have demonstrated an underappreciated role of DDR is to combat MCDD during stem cell migration, and in the absence of fully functioning DDR machinery, stem cells fail to migrate. Bringing our observations together we propose a model where migrating cells go through a “migration-damage-repair-migration” cycle as they move towards the wound site (Figure 4 J). Our findings confirm that migration leads to DNA damage *in vivo* in stem cells and, in the light of earlier findings in cell lines [1–3], may represent an evolutionary conserved cost of this process. Both ageing and oncogenic phenotypes thought to be caused by mutations due to replicative stress may also result from genome instability incurred during cell migration. This could be an under-appreciated source of further genomic heterogeneity in highly invasive cancer cells that encounter tight spaces in the tissue microenvironment [3,34–36]. Future work in planarians on naturally occurring MCDD will help to reveal the regulatory interplay between stem cell migration and DNA-repair processes.

## Funding

This work was supported by the Medical Research Council (Grant number MR/M000133/1), Biotechnology and Biological Sciences Research Council (Grant number BB/K007564/1) awarded to AAA. S.S is funded by the Clarendon Scholarship (University of Oxford) and by the Elizabeth Hannah Jenkinson Fund. N.K. was funded by a Marie Sklodowska Curie individual fellowship by Horizon 2020. A.D. is funded by a BBSRC DTP studentship (BB/J014427/1). M.A.H. and J.M.T. acknowledge funding from the MRC Strategic Partnership Funding (MC-PC-12004) for the CRUK/MRC Oxford Institute for Radiation Oncology.

## Author contributions

A.A.A., SS and PA conceived the study. A.A.A. P.A. and S.S designed the study, analysed the data. A.A.A. and S.S. wrote the manuscript. P.A. optimized the shielded irradiation assay, SS, D.S. and P.A performed experiments. D.S., N.K. and A.D. helped S.S. in the initial characterization and cloning of genes. J.M.T. and M.A.H. helped with use of the irradiation facilities and the design of the shielded irradiation assay. All authors were involved in reviewing and editing the manuscript.

## Declaration of interests

The authors declare no competing interest

## Supplementary figures

**Figure S1.**
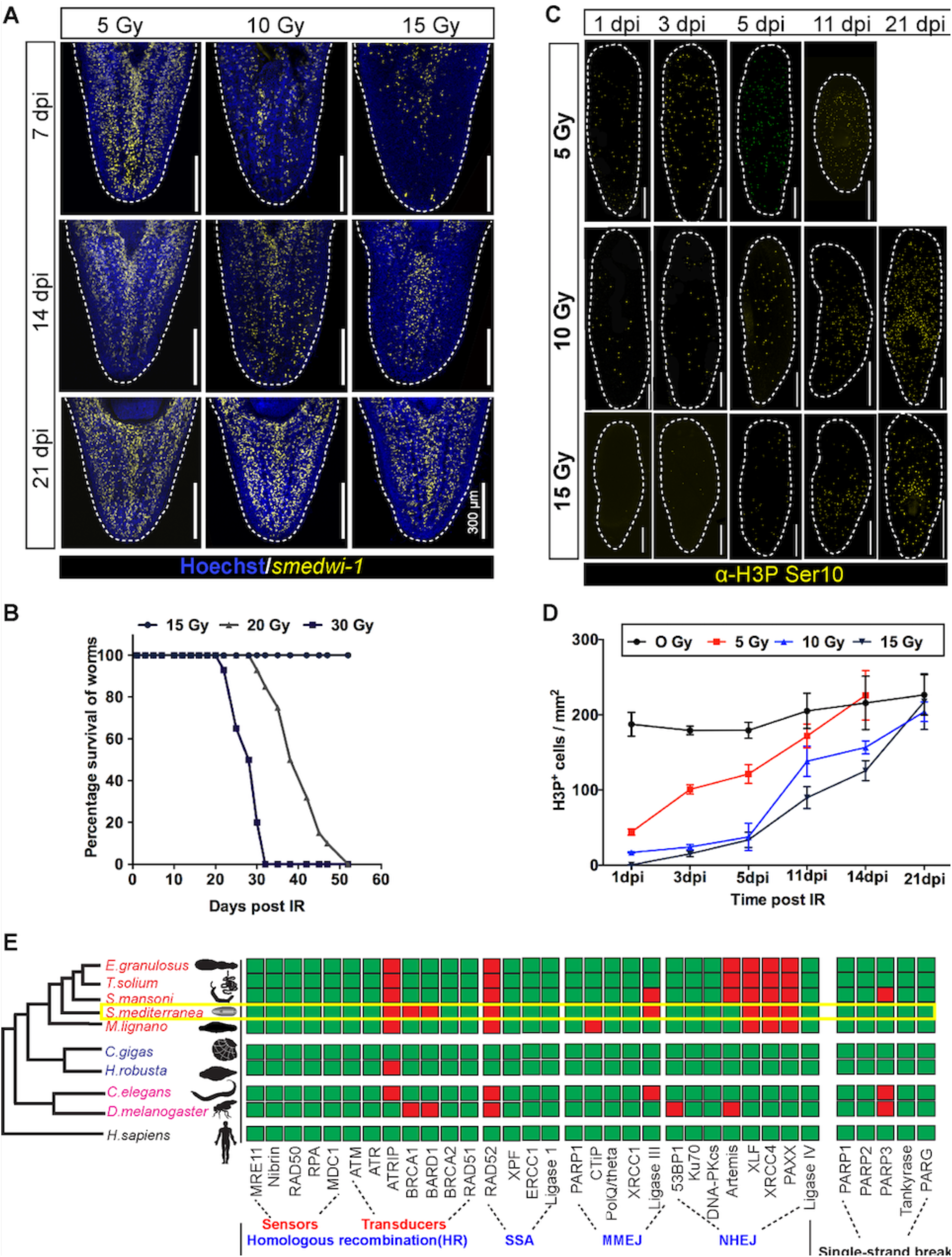
(Related to figure 1) Dynamics of stem cell proliferation and repair kinetics of DNA damage in planarian stem cells. **(A)** FISH showing repopulation of *smedwi-1*^*+*^ stem cells after different doses of IR at indicated days post IR (dpi). Scale bar: 300 μm **(B)** Survival curve indicating the percentage of animals alive after exposure to different doses of ionising radiation, n=15 per dose. Un-irradiated and worms exposed up to 15 Gy IR showed 100 % survival. **(C)** Immunostaining with mitotic marker Anti-H3 phosphorylated-ser10 (H3pSer10) showing the repopulation of mitotic cells (yellow) after exposure to different doses of gamma IR (5,10,15 Gy) at indicated days post IR (dpi). Scale: 500 μm **(D)** Quantification shows the repopulation kinetics of mitotic cells at different doses of irradiation (n = 5 per condition). Results are expressed as mean ± SD. **(E)** Presence (green) or absence (red) of conserved DNA repair genes involved during Double-stranded break repair (DSB) as sensors and transducers in homologous recombination, single stranded annealing (SSA), microhomology-mediated end-joining (MMEJ) and Non-homologous end joining (NHEJ), single stranded break repair in metazoans. The yellow box highlights the DNA repair proteins in *S. mediterranea*.

**Figure S2.**
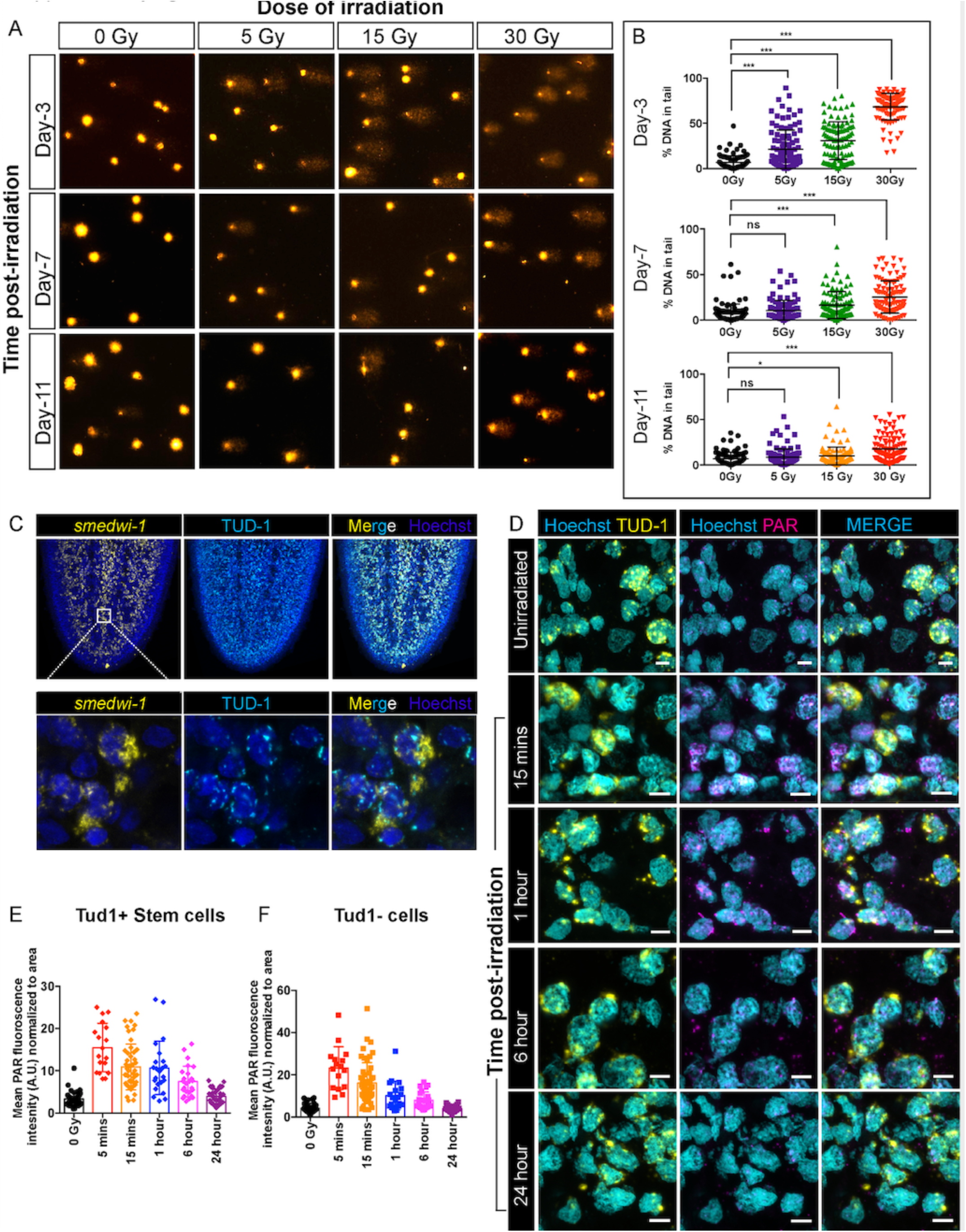
(Related to figure 1) Detection of DNA breaks and DNA damage response in planarian cells after irradiation. **(A)** COMET assay showing the extent of DNA breaks (Comet-shaped tails) on dissociated planarian cells at 3, 7 and 11 days post exposure to 5, 15 and 30 Gy of gamma IR **(B)** Quantification of the percentage of tail DNA in individual comet at indicated time points and dose of IR. (n= 100 comets per condition/time point). **(C)** *Smedwi-1* FISH and Tudor-1 (TUD1) immunostaining showing the presence of perinuclear chromatoid bodies (TUD1+, Cyan) in smedwi-1+ stem cells (Yellow). Nucleus is stained with Hoechst (Blue). **(D)** Double immunostaining with Anti-TUD1(Yellow) and Anti-Poly ADP Ribose (PAR, Magenta) antibodies showing increase in nuclear PAR formation in irradiated planarian cells compared to unirradiated controls; Scale bar: 10 um **(E-F)** Quantification of PAR fluorescence normalised to the area of nucleus from individual TUD1+ stem cells (E) and TUD1-differentiated cells (F) at 5 mins, 15 mins, 1 hour, 6 hour and 24-hour post exposure to 5 Gy IR. The nucleus is stained with Hoechst and pseudo-coloured to cyan. The perinuclear TUD1 staining and Hoechst stained nucleus is used to measure the area of individual nuclei.

**Figure S3.**
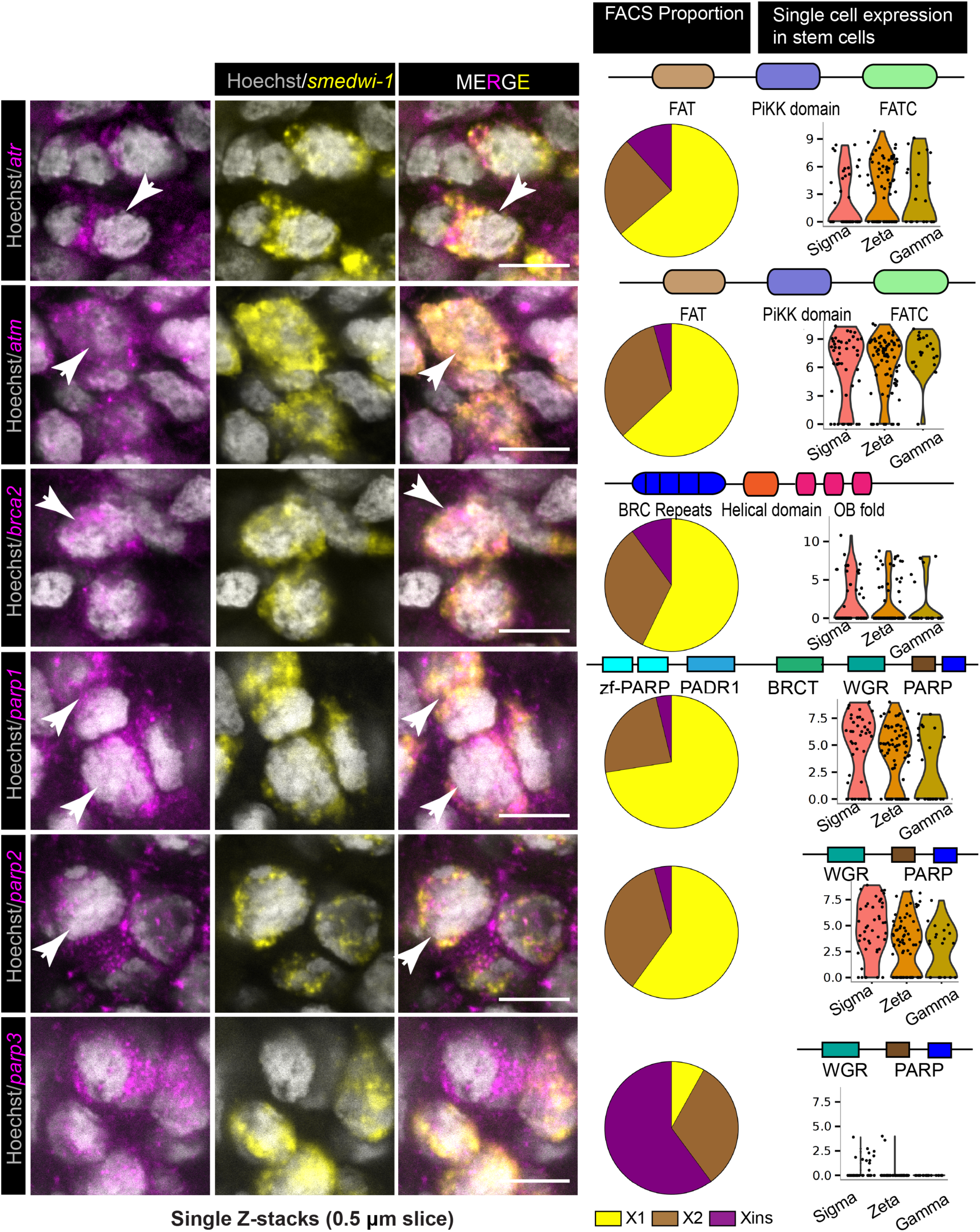
(Related to figure 1) Expression of DNA repair genes in planarian stem cells. Representative double FISH using *smedwi-1* (stem cell marker, yellow) and respective DNA damage response gene (*atr, atm, brca2, parp1, parp2 and parp3*). Arrow mark represents the double positive cells. Scale bar: 5 μm. Proportional expression of the respective DDR genes in FACS sorted planarian cells [Yellow (X1), Brown (X2), Magenta (Xins)] (Dataset from Dattani et. al., 2018). Violin plots showing the single cell expression of the respective DDR genes in sigma, zeta and gamma stem cells. (Dataset from *Wurtzel et al., 2016)* https://radiant.wi.mit.edu/app/. Conserved domains within DDR genes are depicted.

**Figure S4.**
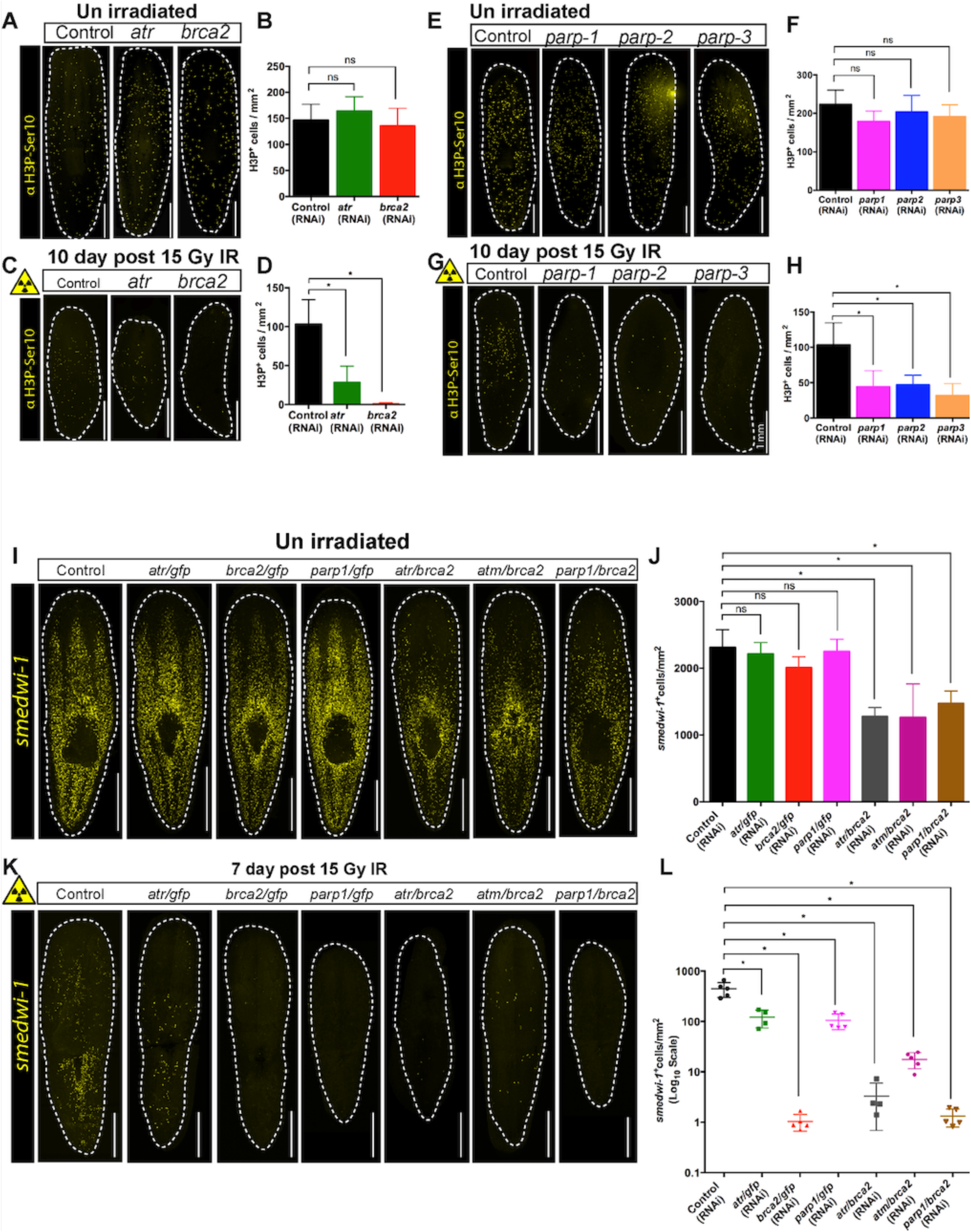
(Related to figure 1) Role of DNA repair gene in planarian stem cell maintenance. **(A)** Immunostaining with H3pSer10 shows the presence of mitotic cells in control worms and after *atr* and *brca2* RNAi. **(B)** Quantification of H3P^+^ cells show no significant difference in cell division after *atr* and *brca2* RNAi. (n=5 per RNAi) Results are expressed as mean ± SD and students’ t test used for analysis (*p<0.05) (ns = not significant). **(C)** Immunostaining with H3pSer10 showing the repopulation of mitotic cells in control worms and after *atr* and *brca2* RNAi worms, 10 days post 15Gy IR. **(D)** Quantification of H3P+ cells show a significant difference in the repopulation of mitotic cells after *atr* and *brca2* RNAi after irradiation, 10 days post 15 Gy IR. (n=5 per RNAi) Results are expressed as mean ± SD and students’ t test used for analysis (*p<0.05) (ns = not significant). **(E)** Immunostaining with H3pSer10 shows the presence of mitotic cells in control worms and after *parp1, parp2* and *parp3* RNAi. **(F)** Quantification of H3P^+^ cells show no significant difference in cell division after *parp1, parp2* and *parp3* RNAi. (n = 5 per RNAi) Results are expressed as mean ± SD and students’ t test used for analysis (*p<0.05) (ns = not significant). **(G)** Immunostaining with H3pSer10 showing the repopulation of mitotic cells in control worms and after *parp1, parp2* and *parp3* RNAi worms, 10 days post 15Gy IR. **(H)** Quantification of H3P+ cells show a significant difference in the repopulation of mitotic cells after *parp1, parp2* and *parp3* RNAi after irradiation. (n=5 per RNAi) Results are expressed as mean ± SD and students’ t test used for analysis (*p<0.05) (ns=not significant). **(I,K)** *smedwi-1* FISH showing the distribution of stem cells after combination of RNAi from different DNA repair genes in unirradiated and after 7 days post 15 Gy IR. GFP dsRNA was used as a control and also added to single RNAi of DDR genes. (J,L) Quantification of *smedwi-1*^*+*^ cells/mm2 in individual worms after RNAi in unirradiated (J) and after irradiation (L). Results are expressed as mean ± SD in log_10_ scale (L) (student’s t-test; *p<0.05).

**Figure S5.**
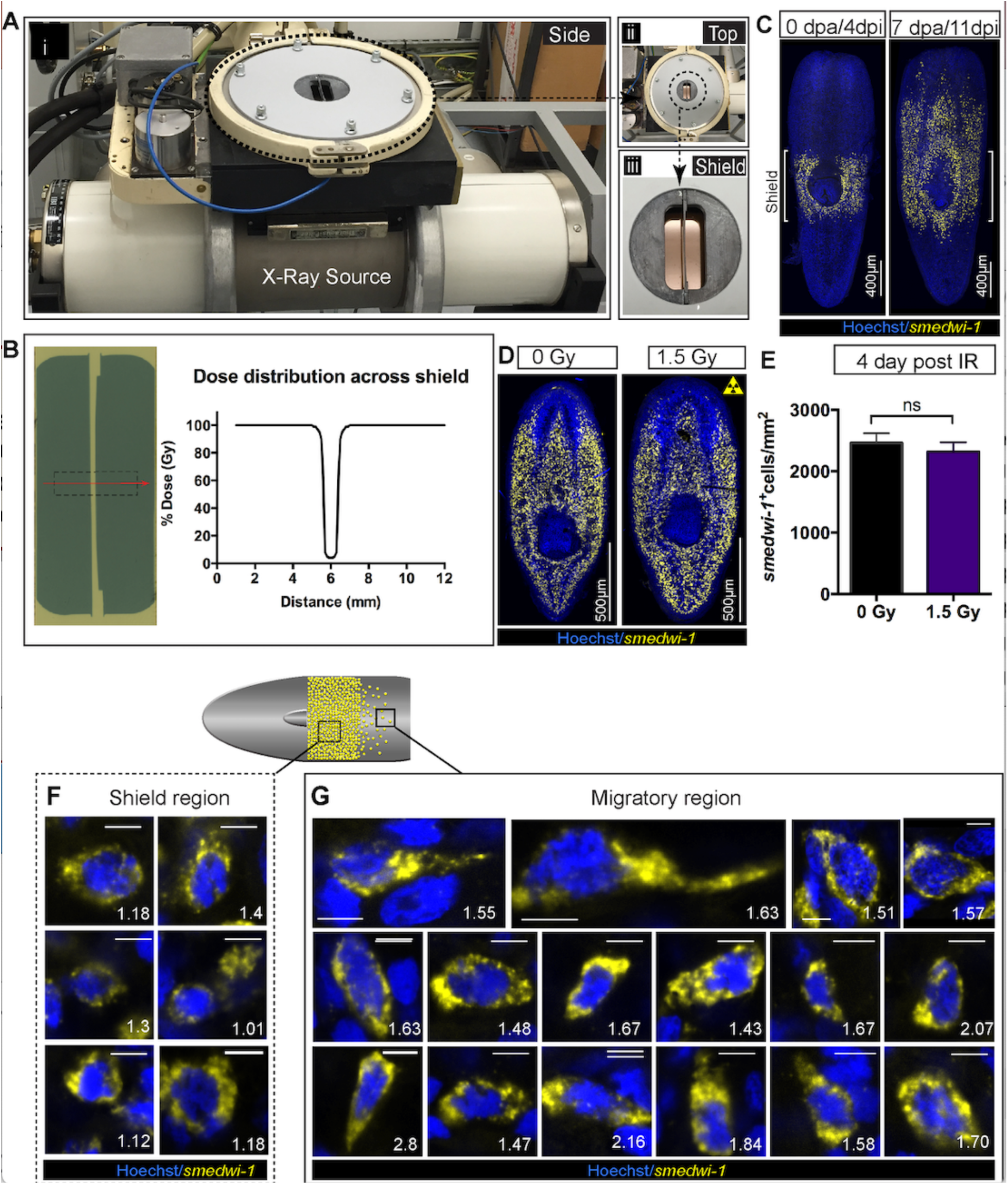
(Related to figure 2) Shielded irradiation assay to study stem cell migration and change in nuclear aspect ratio in migrating cells. **(A)** Experimental set up showing the shielded irradiation assay (i), with a top view showing the position of the shield (ii) and focussing on the shield (iii). **(B)** Dose distribution across the lead strip showing greater than 95% attenuation of X-Ray [4]. An exposure of 30 Gy corresponds to a dose to the cells directly above the shield-protected region of less than 1.5 Gy. **(C)** Representative FISH of *smedwi-1* showing the distribution of stem cells (yellow) after shielded irradiation assay and the migration of stem cells from the shield after 7-day post head amputation. Brackets “[] “represents the position of the shield. Scale bar =400 μm. **(D)** Representative *smedwi-1* FISH showing survival of stem cell after 4-day post 1.5 Gy of IR. **(E)** Graph represents *smedwi-1*^*+*^ cells/mm^2^ showed no significant difference in stem cell maintenance after 1.5 Gy IR. (n =5, *p<0.05, ns=not significant). **(F-G)** Representative FISH showing stem cells (*smedwi-1*^*+*^) with extended protrusions in migratory cells (F) and stationary cells from the shielded region (E). Nuclei stained with Hoechst (blue). Images are shown as single confocal Z-stack (0.32μm). Number represents the NAR value of individual cells, plotted in Fig. 2C. Scale bar = 5 μm.

**Figure S6.**
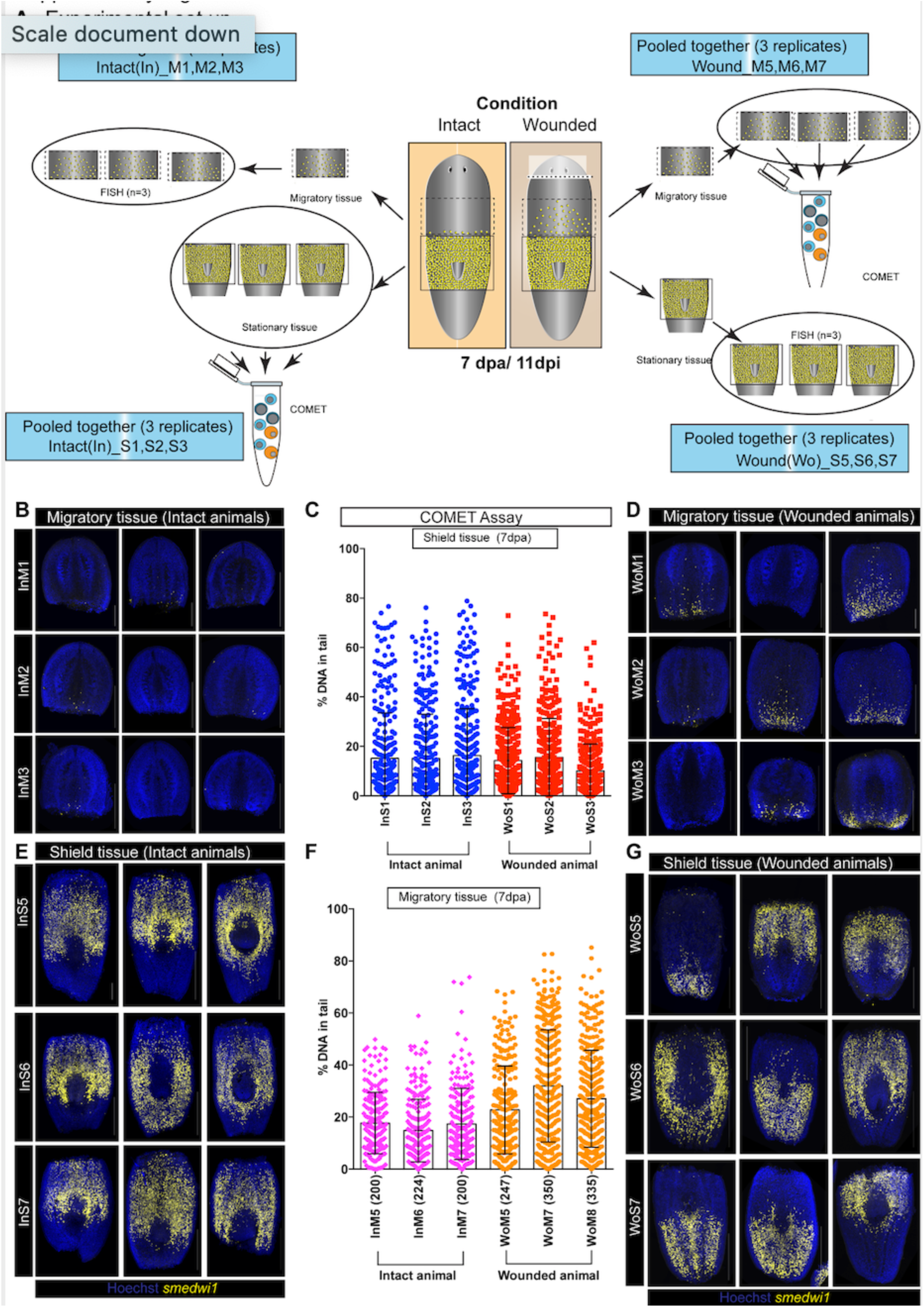
(Related to figure 2) COMET assay to detect DNA breaks during stem cell migration. **(A)** A schematic showing the experimental strategy to perform COMET assay coupled with *smedwi-1* FISH to accurately measure DNA breaks in shielded tissue and migratory tissue. Planarian worms were decapitated (Wounded) 4 days after the shielded irradiation assay and at 7-day post decapitation (7dpa/11dpi) the shielded tissue and the migratory tissue were amputated. Individual migratory tissue fragments from intact and wounded animals were used for COMET assay and the corresponding shielded tissue were used for *smedwi-1* FISH and vice-verse. Each FISH is performed in 3 batches (InM1, InM2, InM3) with 3 worms per batch. The panels are named as “In” or “Wo” corresponding to <u>In</u>tact or <u>Wo</u>unded. “M” and “S” corresponds to <u>M</u>igratory tissue or <u>S</u>hielded tissue. The numbers denote different batch of experiment. The corresponding shielded tissue from these worms were pooled for COMET assay. **(B,D)** *smedwi-1* FISH from migratory tissue of intact animals showing no migration compared to migrating *smedw-1+* stem cells in wounded animals (D). **(C)** Graph represents the percentage of DNA in tail from individual comets from cells in the shielded tissue from intact and wounded animals (300 comets were analysed per condition, n= 900 comets from intact animals and n=900 from wounded animals). **(E, G)** *smedwi-1* FISH from shielded tissue of intact (E) and wounded animals (G) showing the accuracy of cutting the migratory region. The tip of the tail was also amputated before the COMET assay / FISH to reduce the number of dead cells from the irradiated tissue. **(F)** Graph represents the percentage of DNA in tail from individual comets from cells in the migratory tissue from intact and wounded animals showing an increase in DNA breaks in cells from the migratory region compared to the shielded region. Numbers in parenthesis depicts the total number of comets analysed. The cumulative data from F is used in Figure 2L.

**Figure S7.**
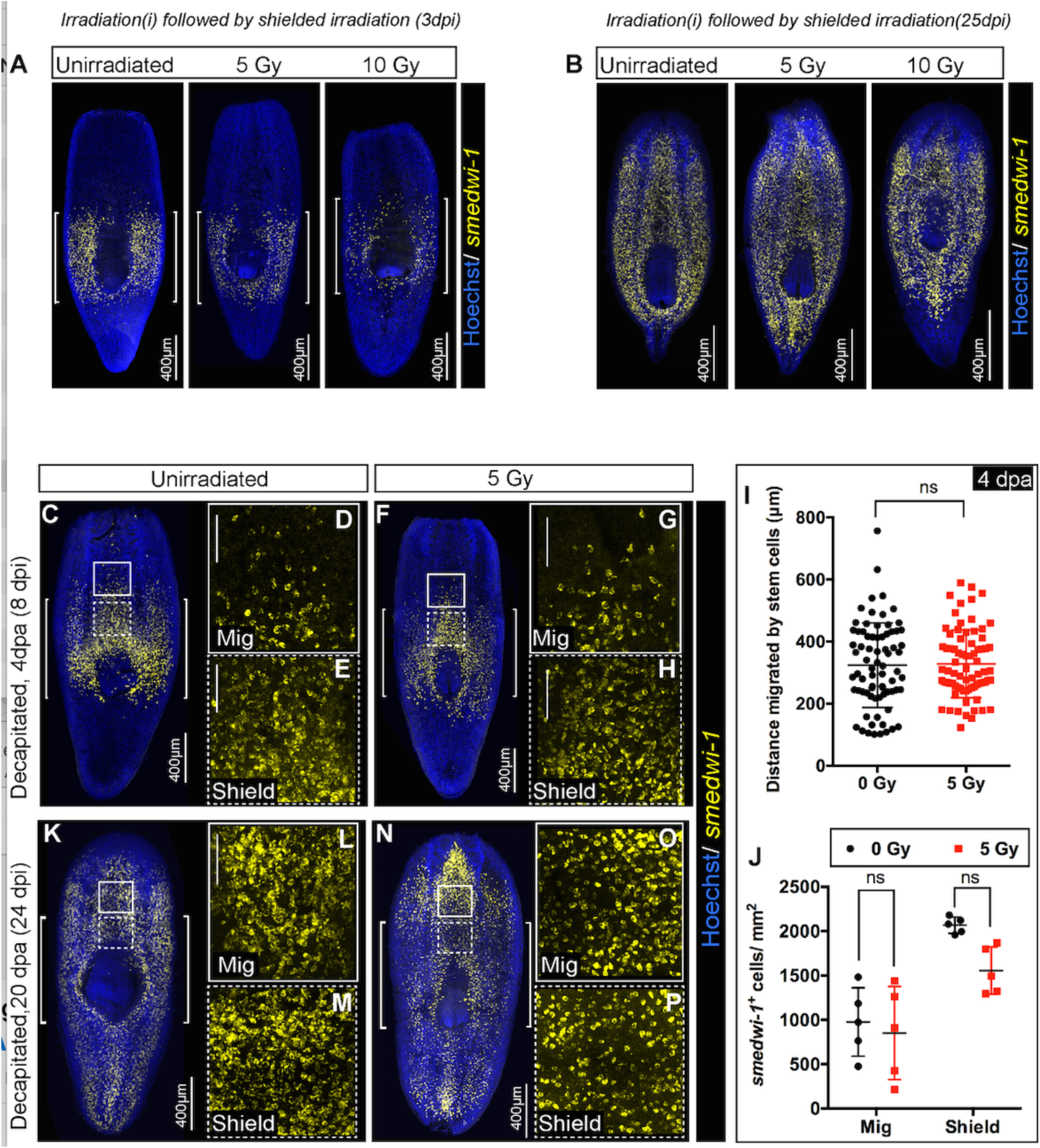
(Related to figure 3) Stem cells pre-loaded with damage before wounding show delays in migration. **(A)** Representative FISH showing the distribution of stem cells in the shield after 5 and 10 Gy of IR followed by the shielded irradiation assay (3day post IR). [] brackets indicate the shielded area. Scale bar: 400 μm. **(B)** The stem cells can eventually migrate and rescue the whole animal. Representative FISH showing the stem cells migrate and reach the wound and rescue the worm (25-day post IR). Scale bar: 400 μm. **(C-P)** Worms are irradiated with a low dose of IR (5 Gy) when cells start to migrate (4dpa, **C-H**) and cells reach the wound (20dpa, **K-P**). Worms are fixed after 24 hours to check the survival of the migratory stem cells **(B-G)** Representative *smedwi-1* FISH showing the sensitivity of migrating cells and cells in the shielded region. The region counted for analysis is marked with a box (bold: migratory region, dotted: Shielded region). [n=5 per condition, Scale bar: 200 μm (B, E), 100 μm (C, D, F, G)]. **(I)** Distance migrated by stem cells suggests no significant sensitivity of stem cells to IR (at least 10 distant cells are counted from individual worms and plotted in the graph. Results are expressed as mean ± SD and students’ t test used for analysis (*p<0.05, ns=not significant) n = 5 worms/condition. **(J)** Quantification of *smedwi-1*^*+*^ cells/mm^2^ cells (yellow) in the shielded region and in the migratory region. Rate of decrease in the migratory field and in the shielded region is plotted in the graph. The region counted for analysis is marked with a box (bold: migratory region F, I, L, O, dotted: Shielded region G, J, M, P). Cells are counted by making a 300 x 300 box anterior to the shield (counted as migratory field) and 300 x 300 box posterior to the shield (counted as shielded region). Results are expressed as mean ± SD and Tukey’s multiple comparison test is used to check for significance. Each dot corresponds to *smedwi-1*^+^cells/mm^2^ in individual worm, n=5, *p<0.05, ns = not significant).

